# Temperate phages as self-replicating weapons in bacterial competition

**DOI:** 10.1101/185751

**Authors:** Xiang-Yi Li, Tim Lachnit, Sebastian Fraune, Thomas C. G. Bosch, Arne Traulsen, Michael Sieber

## Abstract

Microbial communities are accompanied by a diverse array of viruses. Through infections of abundant microbes, these viruses have the potential to mediate competition within the community, effectively weakening competitive interactions and promoting coexistence. This is of particular relevance for host-associated microbial communities, since the diversity of the microbiota has been linked to host health and functioning. Here, we study the interaction between two key members of the microbiota of the freshwater metazoan *Hydra vulgaris*. The two commensal bacteria *Curvibacter* sp. and *Duganella* sp. protect their host from fungal infections, but only if both of them are present. Coexistence of the two bacteria is thus beneficial for *Hydra*. Intriguingly, *Duganella* sp. appears to be the superior competitor *in vitro* due to its higher growth rate when both bacteria are grown seperately, but in coculture the outcome of competition depends on the relative initial abundances of the two species. The presence of an inducible prophage in the *Curvibacter* sp. genome which is able to lytically infect *Duganella* sp., led us to hypothesise that the phage modulates the interaction between these two key members of the *Hydra* microbiota. Using a mathematical model we show that the interplay of the lysogenic life-cycle of the *Curvibacter* phage and the lytic life-cycle on *Duganella* sp. can explain the observed complex competitive interaction between the two bacteria. Our results highlight the importance of taking lysogeny into account for understanding microbe-virus interactions and show the complex role phages can play in promoting coexistence of their bacterial hosts.

## Introduction

Microbial communities are often highly diverse and it is increasingly being recognized that this diversity is key to the ecological functions of these communities [1, 2]. It is thus crucially important to understand the mechanisms allowing for the coexistence of microbial species found in natural communities.

Bacteriophages, the viruses of bacteria, are found across virtually all habitats harbouring microbial communities. Their ubiquity and the observation that globally they are responsible for the turnover of vast amounts of microbial biomass every second [3, 4, 5] has lead to the suggestion that they may play an important role in structuring bacterial communities and maintaining microbial diversity [6]. In particular, theoretical and experimental studies show that they can regulate competitively dominant species or even specific strains via a mechanism termed “kill the winner” [7, 8, 9]. This allows for the coexistence of less competitive species that would otherwise be excluded, thus promoting diversity.

This is in particular relevant for microbes living in and on animal and plant hosts (the microbiota) as there is growing evidence that the composition and functioning of the microbiota and the well-being of the host are closely intertwined. In particular, high microbial diversity has been found to correlate with healthier hosts [10, 11, 12], in line with previous results relating diversity and community functioning [13, 14].

As in many other environments, it has been shown that microbiota are accompanied by an abundant community of viruses [15, 16, 17, 18, 19]. While the majority of previous studies have focused on lytic viruses, recent studies have pointed to the prominent role of lysogeny in microbiota-phage interactions [20, 21] and in fact, an estimated 5% of the human gut bacterial gene content codes for prophage proteins [22]. This suggests that the relationship between phages and bacteria in the microbiota goes beyond predatory or parasitic interactions, and in fact temperate phages have been implicated in increasing the competitive fitness of their lysogenic hosts [23] and driving microbial evolution [24, 25].

Here, we investigate how phages mediate the competition between two key members of the natural microbiota of the freshwater polyp *Hydra vulgaris*. The two bacterial species, *Curvibacter* sp. and *Duganella* sp., are able to protect the polyp against fungal infections and, interestingly, this antifungal activity is strongly synergistic and greatly diminished when one of the species is absent [26]. Coexistence of the two species is thus beneficial for the the host. However, as shown by Li et al. [27] *Duganella* sp. appears to have a much higher growth rate than *Curvibacter* sp. when both are grown in monoculture *in vitro* (Fig. 1a), suggesting that it is competitively dominant. But when both species are grown in coculture at varying initial proportions, the growth rate of *Duganella* shows a strong non-linear dependence on the initial *Curvibacter* sp. frequency. Intriguingly, *Duganella* sp. growth is suppressed at both low and high initial *Curvibacter* sp. frequencies, but not at intermediate frequencies (Fig. 1b).

**Figure 1.**
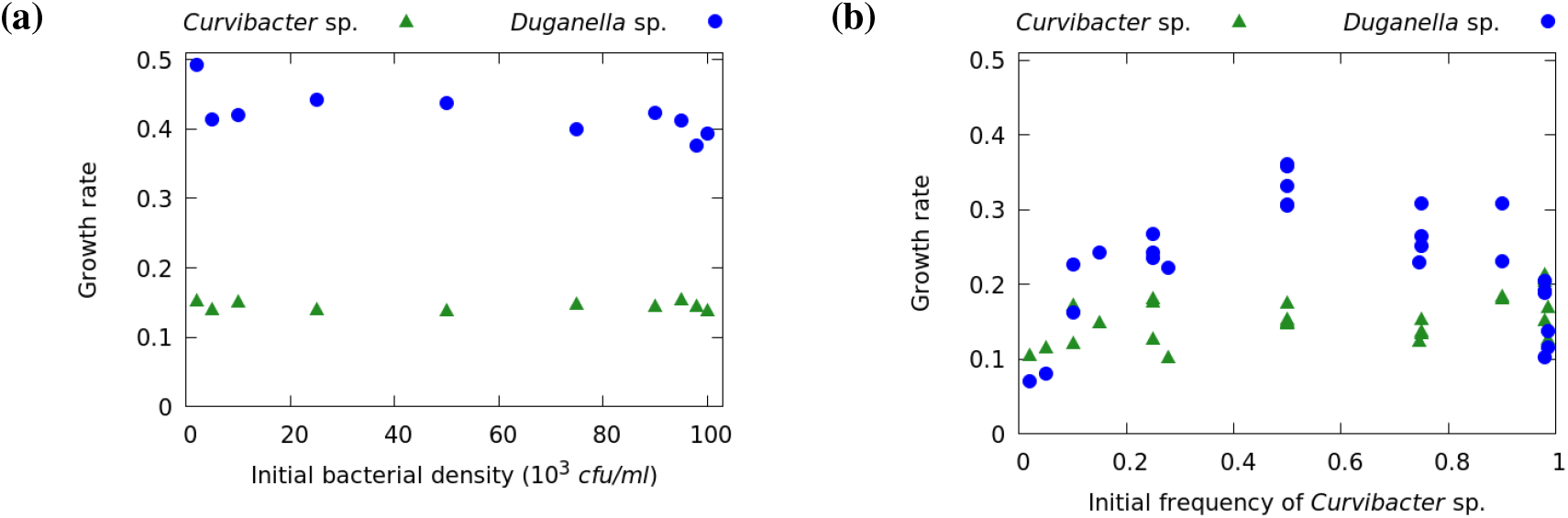
(a) In monoculture the growth rates of *Curvibacter* sp. and *Duganella* sp. do not depend on the initial densities. (b) In coculture the growth rate of *Duganella* sp. changes non-linearly with the initial frequency of *Curvibacter* sp., showing a maximum at intermediate frequencies and inhibition especially at low and high frequencies of *Curvibacter* sp. The growth rate of *Curvibacter* sp. on the other hand does not substantially depend on the initial frequencies. (Adapted from Li et al. [27].)

A recent study has shown that different *Hydra* species harbor a diverse virome and it has been suggested that phages play a role in regulating the *Hydra*-associated microbiota [28, 29]. Indeed we found an intact prophage sequence in the genome of *Curvibacter* sp., which could be reactivated and isolated from bacterial cultures (Fig. 2). Interestingly, the phage can infect and lytically replicate on *Duganella* sp. This leads us to hypothesise that the prophage acts as a self-replicating weapon against *Duganella* sp., thereby playing a key role in modulating the competition between the two bacterial species in the observed non-intuitive way.

**Figure 2.**
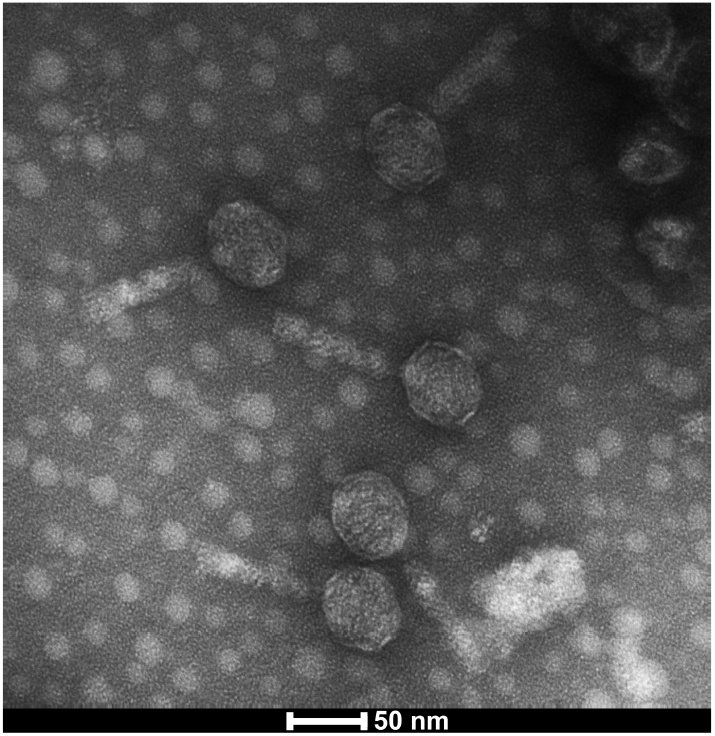
Transmission electron micrograph of mature phages isolated from *Curvibacter* sp. showing the icosahedral head containing the DNA and the tail fibre.

To test this hypothesis, we build a mechanistic bacteria-phage model where, crucially, both the lysogenic cycle on *Curvibacter* sp. and the lytic cycle on *Duganella* sp. contribute to phage population growth. Our model shows that the interplay between the two phage life cycles explains the observed frequency-dependent interactions between the two bacteria and that neither life cycle alone can give rise to the observed competitive interactions.

## Material and Methods

### Isolation of *Curvibacter* phage

*Curvibacter* sp. was grown in R2A broth medium and upon reaching the exponential growth phase mitomycin C was added at a final concentration of 0.05 *μ*g/ml to induce the prophage. After an inoculation time of 16 hours bacterial cells were removed by filtration (pore size 0.2 *μ*m). Bacteriophages were further purified by CsCl density gradient ultracentrifugation [28]. Purified phages were negatively stained in 2% (w/v) aqueous uranyl acetate and visualized by transmission electron microscopy (TEM) using a Tecnai BioTWIN at 80 kV and magnifications of 40,000–100,000X. Infectivity of *Duganella* sp. by *Curvibacter* phage was tested by spot assays according to the protocol described by Adams [30].

### The model

We start by deriving a model of the interactions between the two bacteria and the phage in a well-mixed batch growth culture with a fixed amount of initial nutrients. This allows us to make predictions about the impact of phages on both bacterial species in order to understand the non-linear competition between them *in vitro*.

We denote the densities of the two bacteria (cells/ml) with *C* (*Curvibacter* sp.) and *D (Duganella* sp.) and the density (particles/ml) of free phages with P. In the absence of phages both bacteria take up nutrients *N* (*μ*g/ml) and grow according to Monod kinetics

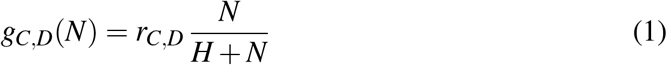

with maximum growth rates *r_C,D_* (h^−1^), respectively. The half-saturation constant *H* (*μ*g/ml) and nutrient conversion efficiency *c* are assumed to be the same for both bacteria.

We incorporate the two distinct phage life cycles as depicted in Fig. 3 in the following way. The lytic life cycle generally involves utilization and killing of the host cell and thus conforms to the classical analogy of phages as predators of their hosts. Consequentely, for phage reproduction via lytic infections of *Duganella* sp. we assume that phage adsorption and infection follow mass action kinetics with adsorption rate *φ* (h^−1^), which leads to lysis and loss of the infected host cell resulting in the release of *β* (cell^−1^) new phages. We include phage production via the lysogenic cycle with the function *d(C*, D), which describes the induction rate of the prophage in *Curvibacter* sp. and subsequent release of new virions via budding through the cell wall, which does not kill the host cell. Note, that this process potentially depends on the densities of both bacteria. The lysogenic cycle is characterized by integration of the phage genetic information into the genome of the host cell, which commonly renders the prophage-carrying host cell immune to infections by related phages [31]. Due to this superinfection inhibition we assume *Curvibacter* sp. to be completely resistant to lytic infections by the phage. Free phages are assumed to decay with a rate *m* (h^−1^).

**Figure 3.**
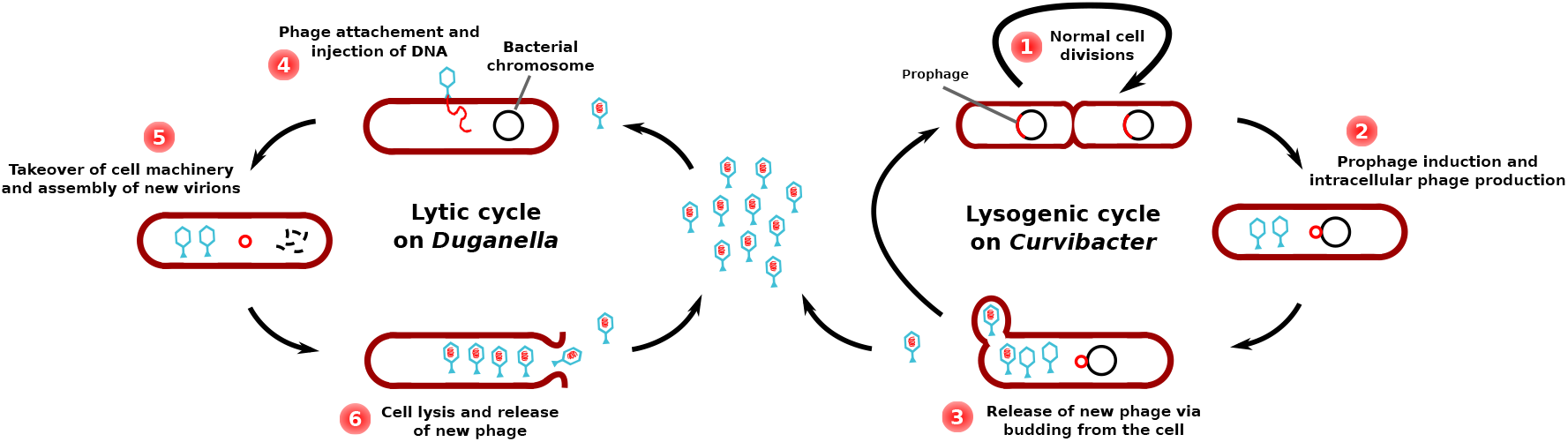
Population growth of free phages is the sum of phage replication via the lytic life cycle on *Duganella* sp. (left) and the lysogenic life cycle on *Curvibacter* sp. (right). The lysogenic cycle is characterized by the integration of the phage DNA into the *Curvibacter* sp. chromosome as a prophage. Cells divide normally (1) and induction of the prophage leads to the production of virions (2). Mature phages are released by non-lethal extrusion or budding (3). The lytic cycle is initiated by infection of a *Duganella* sp. cell by a free phage (4), followed by phage replication (5) and release of the new phages (6). The last step results in the lysis and death of the *Duganella* host cell.

With this the model describing the dynamics of nutrients, bacteria and phages reads

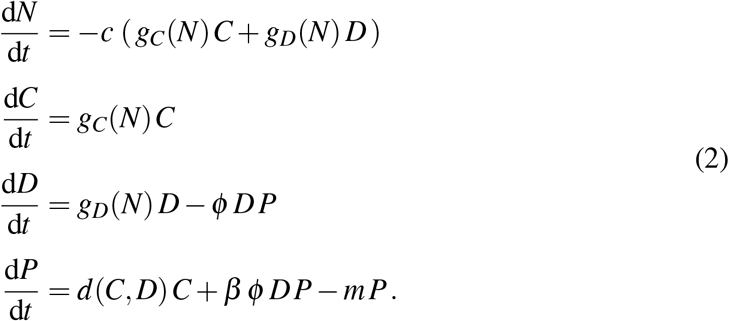

The key feature of this model is that phage population growth is the sum of lytic reproduction on *Duganella* sp. and lysogenic phage production from *Curvibacter* sp.

So far, we have not explicitly defined the prophage induction rate *d(C, D)*. Although the induction of lysogenic phages can have a substantial impact on bacterial population dynamics, the factors that induce prophages and modulate the rate of induction are still largely unknown [32]. Some of the known factors that can activate prophages include the host’s SOS response following cell damage, external triggers, and spontaneous induction by stochastic gene expression [33]. Since our results do not depend on the particular mechanism modulating the induction rate, in the following section we aim to deduce the mathematically simplest form of *d(C*, D) that is consistent with the experimental data.

### Approximating the prophage induction rate

First we need to clarify how the induction rate *d*(*C, D*) influences the *Duganella* sp. growth rate. If nutrients are not limiting, i.e. during exponential growth, we have *g_D_*(*N*) ≈ *rD* and thus the *Duganella* sp. growth rate simplifies to

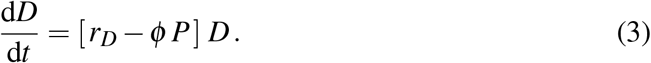

The growth rate is the difference between the *Duganella* sp. growth rate in mono-culture and the losses imposed by the phage. We approximate those losses by assuming that phage dynamics are much faster than bacterial dynamics, implying that the phage density quickly reaches its asymptotic value *P*\* for which *dP/dt* = 0, where the bacterial densities *C* and *D* are essentially free parameters. Solving for this equilibrium value and inserting it into the growth rate (3) of *Duganella* sp. yields

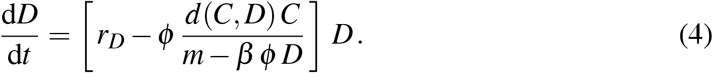

The term in the brackets corresponds to the exponential growth rate of *Duganella* sp. in the presence of *Curvibacter* sp., where the indirect effect of *Curvibacter* sp. is mediated by the phage. Denoting this term with *G* and rewriting it using the frequency *f* = *C/B* of *Curvibacter* sp. in the total bacterial population *B* = *C* + *D* gives

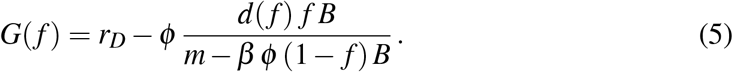

Note that *G*(0) = *r_D_* recovers the growth rate of *Duganella* sp. in mono-culture.

Now we can ask: what is the simplest form of the prophage induction rate *d*(*f*) that would lead to a hump-shaped growth rate *G*(*f*) of *Duganella* sp. as observed in the experiments (Fig. 1b)? The condition that *G*(*f*) attains its maximum at some intermediate *Curvibacter* sp. frequency implies that

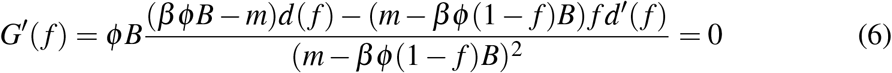

for some 0 < *f* < 1.

Now, if prophage induction rate is independent of *Curvibacter* sp. frequency, we have *d*(*f*) = *p* with some constant *p* > 0. However, in this case *d^f^*(*f*) = 0 and thus

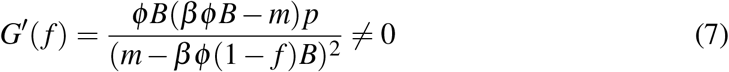

for all *f*. This shows that a constant induction rate does not allow for a hump-shaped growth rate of *Duganella* sp.

A slightly more complex case is given by a linearly increasing induction rate, namely *d*(*f*) = *pf*, with rate constant *p* (particles h^−1^ cell^−1^). In this case *G*′(*f*) = 0 has two solutions, namely *f* = 0 and

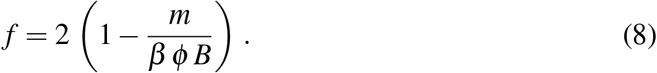

Thus, under the assumption of fast phage and slow bacteria dynamics, a hump-shaped growth rate of *Duganella* sp. is possible if prophage induction rate increases linearly with *Curvibacter* sp. frequency. Accordingly, in the following analysis of the model, we set *d* (*f*) = *pf*.

## Results

For a wide range of parameter values and initial conditions (*N*_0_,*C*_0_,*D*_0_ > 0 and *P*_0_ ≥ 0) the population dynamics described by equations (2) result in the following general pattern (see the Suppl. Fig. 1, for an example): after inoculation of the coculture when nutrients are not limiting, both bacteria grow exponentially. *Curvibacter* sp. keeps growing until nutrients near depletion (*N* → 0), at which point its growth rate approaches zero and its density reaches a final value *C*\*, which depends on the initial density *D*_0_ of *Duganella* sp. and the initial amount *N*_0_ of nutrients. *Duganella* sp. on the other hand will initially also grow exponentially, but in addition to the nutrient levels its population growth is affected by the losses imposed by phage infections.

To avoid the confounding effects of nutrient depletion and to keep the model close to the experimental setup, we restrict our analysis to the exponential growth phase when nutrients are not yet limiting. We define the end of this phase as the time when *Curvibacter* sp. growth rate falls below a certain threshold *ε* > 0, since *Curvibacter* sp. growth rate is only influenced by the current nutrient level. In particular, we numerically calculate the *Duganella* sp. growth rate during the exponential phase as

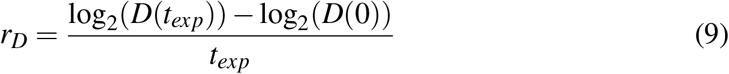

over the time interval [0, *t_exp_*] for which

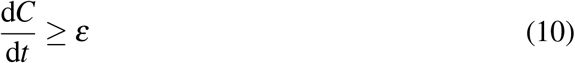

holds. In exactly the same manner we obtain the growth rate *r_C_* of *Curvibacter* sp.

During this time interval the phage is the only factor influencing *Duganella* sp. growth rate and the question is whether the growth dynamics of the phage alone can give rise to the observed non-linear interaction between the two bacteria. To address this question it is important to note that the growth of the phage population is determined by both *Curvibacter* sp. and *Duganella* sp. densities via prophage induction and lytic reproduction, respectively. As we will now show, the contributions of these two pathways to phage growth behave very differently as the relative initial abundances of the two bacteria change.

### Contribution of the lytic cycle

The lytic life cycle is known to be especially effective at high host densities, when each phage particle quickly finds a new host. In this case even a small amount of phage can initiate rapid replication resembling a chain reaction, leading to a quick and notable decline or even collapse of the bacterial host population. If the host density is too low, however, the losses through phage decay are not outweighed by reproduction, the phage can thus not invade the host population and is eventually lost from the system. In our system the so called “replication threshold” [34, 35, 36] - i.e., the minimum host density that allows positive phage growth - can be calculated explicitly by observing that, if there is only lytic reproduction (*d*( *f*) = 0 for all *f*), we have

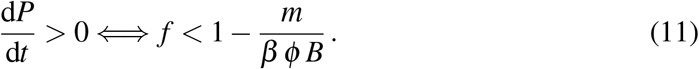

This implies that, with lytic reproduction alone, the phage can grow and have a substantial impact on *Duganella* sp. growth only if the initial *Curvibacter* sp. frequency *f* is relatively low. Conversely, as long as the relative abundance of *Curvibacter* sp. in the total population is too high, the phage does not find enough suitable *Duganella* sp. hosts and can not persist. Note, that the replication threshold increases with total bacterial population density *B* and approaches 1 for very high densities, implying that it is most relevant in the early stages of exponential growth when bacterial densities are still low, which is exactly the case we consider here. In particular, because *Duganella* sp. is growing faster than *Curvibacter* sp. when it is rare, it will eventually reach the replication threshold of the phage, but it does so too late for the phage to grow to substantial densities before nutrients are depleted.

See Fig. 4a for a numerical example of this frequency-dependent impact of lytic phage growth on the two bacteria, which shows the bacterial growth rates as defined above for different initial frequencies of *Curvibacter* sp. and *Duganella* sp. in the case of no lysogenic production. Note, that *r_C_* is constant for all initial *Curvibacter* sp. frequencies, reflecting that during the exponential growth phase *Curvibacter* sp. growth rate is inde-pendent of both the phage and the presence of *Duganella* sp.

**Figure 4.**
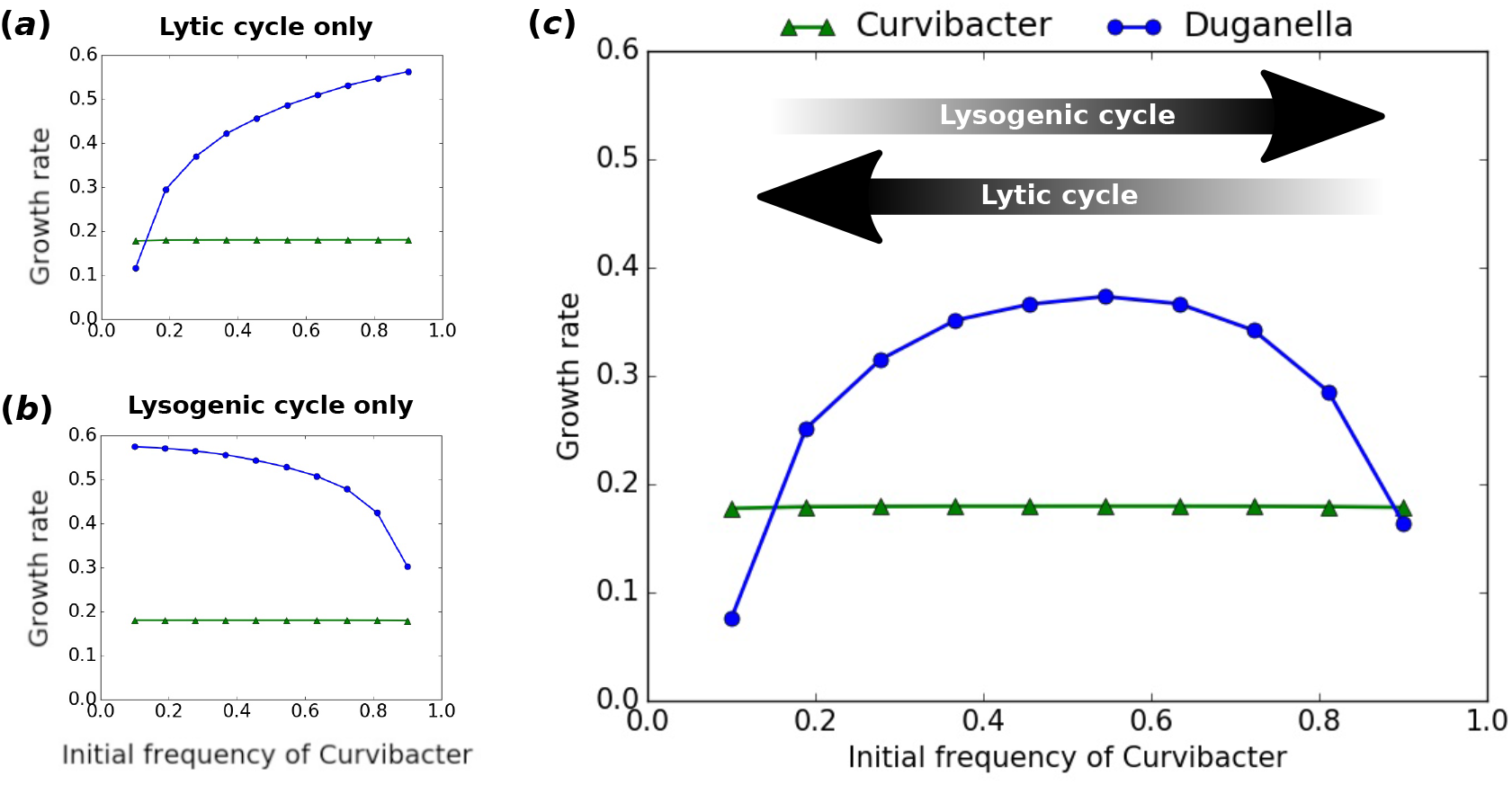
Growth rates obtained from the model during the exponential growth phase of *Curvibacter* sp. and *Duganella* sp. for different initial frequencies of *Curvibacter* sp. (a,b) The growth rates if only the lytic (a) or lysogenic (b) phage life cycle is taken into account. (c) Bacterial growth rates if both life cycles are taken into account. The arrows indicate at which ends of the initial frequency spectrum the two different phage life cycles are most effective. Parameter values for this example: *N*_0_ = 100, *B*_0_ = 10^5^, *P*_0_ = 3.5 × 10^4^, *c* = 10^−5^, *r_C_* = 0.125, *r_D_* = 0.4, *H* = 0.05, *φ* = 10^−8^, *β* = 50, *m* = 0.25, *p* = 0.05.

### Contribution of the lysogenic cycle

Let us now turn to phage reproduction via the lysogenic pathway (denoted by *d* (*C, D*) in Eqn (2)). It is clear that the contribution of the lysogenic pathway to phage growth increases as the frequency of *Curvibacter* sp. in the total population increases. However, lysogenic production alone is generally not enough to sustain the phage at high densities and lytic reproduction is still necessary for the phage to reach densities at which it has a visible impact on *Duganella* sp. growth. We can illustrate how the contribution of the lysogenic pathway increases with *Curvibacter* sp. frequency by setting the density of initially present phages *P*_0_ to zero. In this case the founder population of phages is introduced solely via induction of the prophage from *Curvibacter* sp., emphasizing the contribution of the lysogenic pathway.

In this scenario, if the initial relative abundance of *Curvibacter* sp. is too low, the production of phages via prophage induction is not sufficient to provide a large enough phage population for it to have a substantial impact on *Duganella* sp. before nutrients are depleted. Only when the initial frequency of *Curvibacter* sp. is sufficiently high does the lysogenic cycle produce enough phages to suppress *Duganella* sp. growth (Fig. 4b).

### Combined effects of the lytic and lysogenic cycles

Now, if both pathways are active and phage population growth is the sum of the lysogenic and lytic life cycles, the two patterns described above combine to give the observed nonlinear dependence of the *Duganella* sp. growth rate on the initial composition of the bacterial population (Fig. 4c). Essentially, the two effects appear superimposed on each other, with the *Duganella* sp. growth rate being supressed at both low and high initial *Curvibacter* sp. frequencies.

## Discussion

In this study, we considered how temperate phages affect the interactions between bacteria in a frequency-dependent manner. We developed a mathematical model that takes into account both the lysogenic and the lytic life cycles of temperate phages. Crucially, our model shows that the two pathways show very different efficiencies as host population composition changes.

At low frequencies of the prophage-carrying *Curvibacter* sp., the main route of phage production is via the lytic pathway, resulting in a significant decrease of *Duganella* sp. growth rate (Fig. 4). In this case, the phage has a high chance of encountering a susceptible *Duganella* sp. cell, which allows the phage to spread rapidly and lyse a great portion of the *Duganella* sp. population in the process. The mechanism provides an efficient way for *Curvibacter* sp. to bounce back from very low frequencies, thus preventing extinction and preserving diversity. A similar use of temperate phages has been shown to confer a competitive advantage on prophage carriers in bacterial competition [37], in particular allowing rare invaders to spread and persist in susceptible resident populations [38,39]. The effectiveness of temperate phages as either a persistence or invasion strategy relies on the fact that the lytic pathway is particularity efficient when the resistant prophage-carriers are rare and susceptible hosts are abundant, allowing the rapid spread of the phage as a self-replicating weapon. This is in contrast to antimicrobial toxins, which are most effective when the toxin producers are at high densities [38].

At high relative abundances of *Curvibacter* sp. on the other hand, the lytic pathway is very ineffective as the majority of the bacterial population are resistant lysogens. In this case lytic replication alone is not able to sustain the phage population, and thus without the lysogenic production from *Curvibacter* sp. the phage would be lost from the system. But through increased lysogenic reproduction at high *Curvibacter* sp. densities the phage is not only able to persist, but in fact it can lyse a significant portion of the *Duganella* sp. population and thus diminish its growth rate (Fig. 4).

At intermediate *Curvibacter* sp. frequencies, however, neither life cycle is particu-larily effective, which allows *Duganella* sp. to grow relatively unaffected by the phage. Taken together, this gives rise to the observed hump-shaped dependence of *Duganella* sp. population growth on the initial relative abundance of *Curvibacter* sp. Crucially, our results also show that only by taking both life cycles into account can the observed growth rates be recovered (Fig. 4).

We deduced that a prophage induction rate that is linearly increasing with *Curvibacter* sp. frequency is consistent with the experimental data. Such a concerted prophage induction is for example possible via quorum-sensing mechanisms and this has been hypothesised to allow phages to sense favorable conditions [40]. However, we want to emphasise that our results do not depend on the precise mechanism of prophage induction. In particular, the induction rate does not necessarily have to be linearly increasing and we do not expect prophage induction rates to follow a linear function in nature. Rather, our aim was to show in a “proof-of-principle” manner an example of minimal complexity that gives rise to the observed bacterial growth dynamics.

Our result provides a potential mechanism behind the finding that weak competitive interactions are the dominant type of interactions within host-associated microbial communities [41]. In line with the “killing the winner” mechanism [42] the phage imposes a top-down control of the otherwise stronger competitor *Duganella* sp., potentially alleviating the competitive interactions between the two bacterial species and allowing them to coexist within the microbiota of *Hydra vulgaris*.

While there is no indication that the phage lysogenises *Duganella* sp., there is a possibility that over a longer time span *Duganella* sp. becomes resistant to the phage via lysogenization. This would imply a shift from its short term use as a weapon to a more classical parasitic life style, a shift that has been reported to occur after a few days of competition between phage-free strains of *Escherichia coli* and strains lysogenized by the temperate phage λ [43]. Over longer time spans antagonistic coevolution between bacteria and phages will also likely contribute to bacterial resistance in the microbiota [44], further adding to the complexity of bacterial competition mediated by phages.

The observation that *in vivo* the protection of the *Hydra* host against fungal infections is mediated by the coexistence of both bacterial species [26] suggests, that an active regulation of the induction of the *Curvibacter* sp. prophage could be beneficial for *Hydra*. And, intriguingly, it has recently been shown that *Hydra vulgaris* is able to modify the quorum-sensing signals of *Curvibacter*, thereby changing the phenotype of a bacterial colonizer [45]. This is an example of a mechanism by which a host could potentially manipulate bacterial gene expression, including prophage induction. More generally, the growing recognition of phages as regulators of the microbiota [46, 47] leads to the interesting hypothesis, that the ability to manipulate the induction of prophages to promote the coexistence of synergistic bacteria is itself an evolvable host trait.

In conclusion, we found that, by taking the lysogenic and lytic life cycles of temperate phages into account, a minimal model is able to explain the observed frequency-dependent outcome of competition between two key members of the *Hydra vuglaris* microbiota. Our study elucidates the inherently complex effects of temperate phages on bacterial competition and population dynamics, which are shown to be highly dependent on the composition of the host population. Future studies are aimed at identifying the precise mechanisms of *Curvibacter* prophage regulation and the quantification of its induction rate, which will shed further light on the role of temperate phages in host-associated microbial communities.

## Data Accessibility

The growth rate data shown in Fig. 1 is available in the Data Supplement of Li et al. [27].

## Authors’ contributions

X. L. and M. S. developed and analysed the mathematical model. T. L. carried out laboratory work and obtained microscopy images. X. L, T. L. and M. S. conceived the study. All authors designed the study, contributed to writing the manuscript and gave final approval for publication.

---

**Suppl. Fig. 1.**
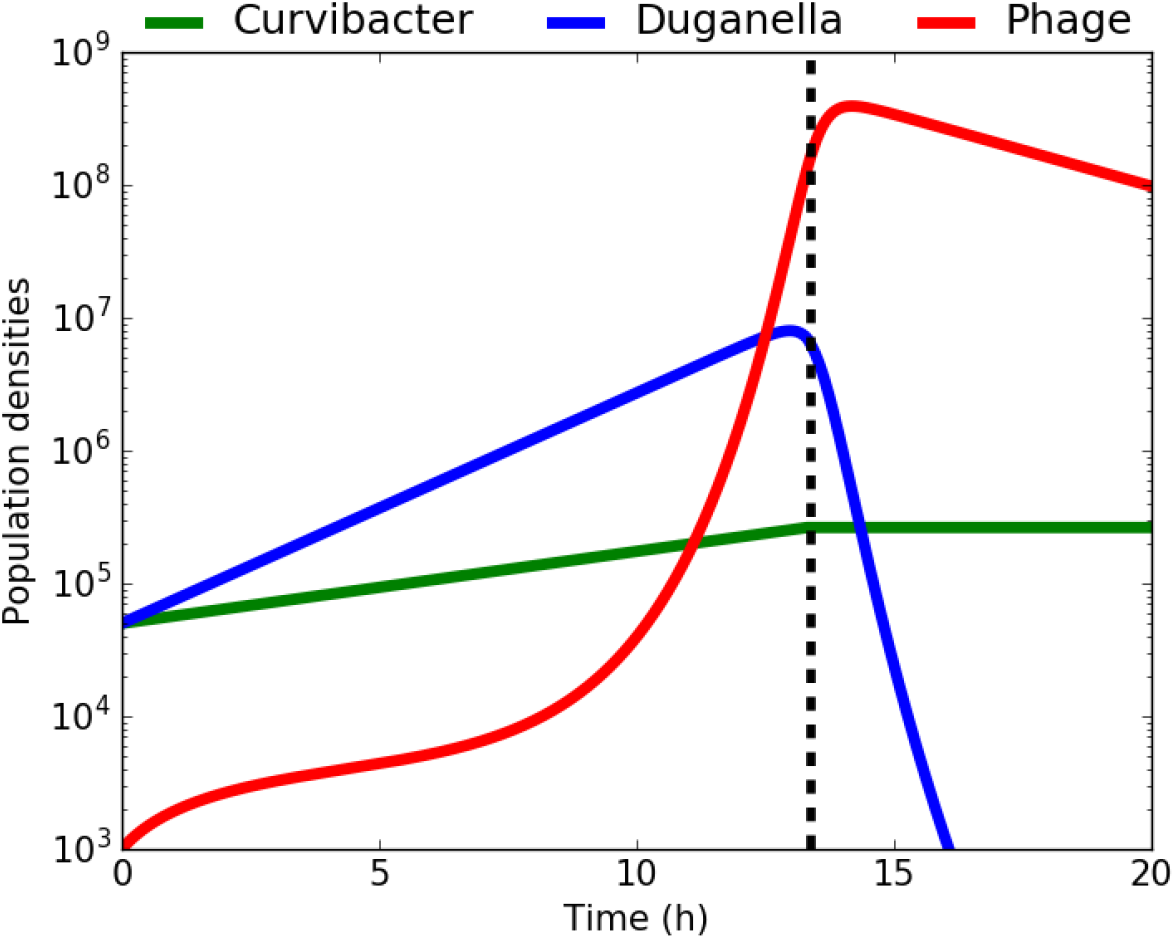
Typical population dynamics obtained from the model (2). Both bacteria initially grow exponentially, until the *Duganella* sp. growth declines due to lytic phage infections and the *Curvibacter* sp. growth slows down due to depletion of nutrients. The vertical dashed line at *t = texp* indicates the end of the exponential growth phase, defined as the time when the *Curvibacter* sp. growth ceases. The *Duganella* sp. density approaches 0 after about 20 hours due to continued phage infections. Parameter values for this example: *N*_0_ = 100, *B*_0_ = 10^5^, *f*_0_ = 0.5, *P*_0_ = 10^3^, *c* = 10^−5^, *r_C_* = 0.125, *r_D_* = 0.4, *H* = 0.05, *φ* = 10^−8^, *β* = 50, *m* = 0.25, *p* = 0.05.

